# Reduced SV2A and GABA_A_ receptor levels in the brains of type 2 diabetic rats revealed by [^18^F]SDM-8 and [^18^F]flumazenil PET

**DOI:** 10.1101/2023.09.10.557061

**Authors:** Yanyan Kong, Fang Xie, Xiuzhe Wang, Chuantao Zuo, Kuangyu Shi, Axel Rominger, Qi Huang, Jianfei Xiao, Donglang Jiang, Yihui Guan, Ruiqing Ni

## Abstract

**Purpose:** Type 2 diabetes mellitus (T2DM) is associated with a greater risk of Alzheimer’s disease. Synaptic impairment and protein aggregates have been reported in the brains of T2DM models. Here, we assessed whether neurodegenerative changes in synaptic vesicle 2A (SV2A), γ;-aminobutyric acid type A (GABA_A_) receptor, amyloid-β, tau and receptor for advanced glycosylation end product (RAGE) can be detected in vivo in T2DM rats.

**Methods:** Positron emission tomography (PET) using [^18^F]SDM-8 (SV2A), [^18^F]flumazenil (GABA_A_ receptor), [^18^F]florbetapir (amyloid-β), [^18^F]PM-PBB3 (tau), and [^18^F]FPS-ZM1 (RAGE) was carried out in 12-month-old diabetic Zucker diabetic fatty (ZDF) and Sprague□Dawley (SD) rats. Proteomic profiling and pathway analysis of the hippocampus of ZDF and SD rats were performed.

**Results:** Reduced cortical [^18^F]SDM-8 and cortical and hippocampal [^18^F]flumazenil uptake were observed in 12-month-old ZDF rats compared to SD rats. [^18^F]florbetapir and [^18^F]PM-PBB3 uptake were comparable in the brains of 12-month-old ZDF rats and SD rats.

**Conclusion:** The findings provide in vivo evidence for regional reductions in SV2A and GABA_A_ receptor levels in the brains of aged T2DM ZDF rats.

## Introduction

T2DM is a major public health concern associated with a greater risk of brain damage, cognitive decline and susceptibility to AD [1]. Decreased hippocampal volume and long-term potentiation and changes in glucose metabolism have been observed in T2DM patients [2]. Postmortem studies in the brains of patients with T2DM have demonstrated loss of synaptic protein and neurotransmitters, including γ-aminobutyric acid type A (GABA_A_) [3]. Studies in T2DM mouse and rat models have demonstrated metabolic alterations, GABA_A_ receptor alterations [4], impairment in memory and learning, and hippocampal synaptic reorganization [5-7]. GABA is the main inhibitory neurotransmitter released at GABAergic synapses and plays an important role in regulating brain activity [3]. GABAergic impairment has been a therapeutic target in T2DM with ongoing clinical trials. The beneficial effect of upregulating GABA_A_ receptors was demonstrated in T2DM models [4]. In addition, advanced glycation end products (AGEs)/RAGE signaling plays an important role in T2DM and is involved in multiple pathways [8]. In addition, increased levels of soluble/insoluble amyloid-beta (Aβ), Aβ plaques, phosphorylated tau, and tau deposits, which are pathological features in the brains of AD patients and animal models, have been shown in the brains of T2DM rats [9-11].

PET imaging for Aβ and tau aggregates has been utilized in AD research and clinical applications to noninvasively map the regional distribution in the brain, such as [^18^F]florbetapir, [^18^F]florbetaben, [^18^F]flutemetamol, and [^11^C]PIB for visualization of Aβ deposits and [^18^F]PI-2620, [^18^F]flortaucipir [12-17], [^18^F]MK-6240, [^18^F]THK-5351, [^18^F]RO948, [^18^F]PM-PBB3, and [^11^C]PBB3 [18-22] for the detection of tau. SV2A imaging using [^18^F]SDM-8 ([^18^F]SynVesT-1), [^18^F]SDM-16, [^18^F]UCB-H, and [^11^C]UCB-J has been developed and has shown reduced synaptic density in AD patients and amyloidosis models [23-27]. For mapping GABA_A_ receptors, [^11^C]flumazenil is the most widely used tracer in both animal models and patients with various diseases [28]. Decreased cerebral uptake of [^11^C]flumazenil has been demonstrated in AD patients and P301L tau mouse models [29]. The development of ligands for RAGE has been challenging. Several tracers that detect the extracellular domain of RAGE have been developed: [^18^F]FPS-ZM1 ([^18^F]RAGER), [^11^C]FPS-ZM1, [^18^F]S100A4, [^99m^Tc]anti-RAGE-F(ab′)_2_, [^64^Cu]Rho-G4-CML, [^64^Cu]MMIA-CML, and [^64^Cu]MMIA-HAS nanoparticles [30-35].

Here, we aimed to assess the levels of synaptic vesicle 2A (SV2A), GABA_A_ receptor, Aβ, tau, and RAGE by in vivo imaging in a widely used T2DM model. Multitracer PET using [^18^F]SDM-8 (SV2A) [25], [^18^F]flumazenil (GABA_A_ receptor) [28], [^18^F]florbetapir (Aβ), [^18^F]PM-PBB3 (tau), and [^18^F]FPS-ZM1 (RAGE) [30] was conducted in aged diabetic ZDF and SD rats, followed by proteomics and pathway analysis of hippocampal tissue.

## Material and Methods

### Animal models

The animal models, including 12-month-old ZDF rats, 12-month-old Sprague□Dawley (SD) rats, and 1.5-month-old C57BL/6J mice, were used in the study (**Table 1**). Rats and mice were housed in ventilated cages inside a temperature-controlled room under a 12-h dark/light cycle. Pelleted food and water were provided ad libitum. Paper tissue and shelters were placed in cages for environmental enrichment. The in vivo PET imaging and experimental protocol was approved by the Institutional Animal Care and Ethics Committee of Fudan University and performed in accordance with the National Research Council’s Guide for the Care and Use of Laboratory Animals and the ARRIVE guidelines 2.0,

**Table 1.** Information on the rats and mice used in the study.

| **Rat** | **Sex** | **Age**<br>**(month)** | **[<sup>18</sup>F]SDM-**<br>**8** | **[<sup>18</sup>F]**<br>**flumazenil** | **[<sup>18</sup>F]FPS-**<br>**ZM1** | **[<sup>18</sup>F]florbetap**<br>**ir** | **[<sup>18</sup>F]PM-**<br>**PBB3** |
| --- | --- | --- | --- | --- | --- | --- | --- |
| SD | M | 12 | 4 | 3 | 1 | 8 | 4 |
| ZDF | M | 12 | 5 | 3 | 5 | 15 | 7 |
| **Mouse** | **Sex** | **Age**<br>**(month)** | **Biodistrib**<br>**ution** | **Blood**<br>**sampling** |  |  |  |
| C57BL/6J | M | 1.5 | 15 | 3 |  |  |  |
| C57BL/6J | F | 1.5 | 15 | 3 |  |  |  |

M, male; F, female mice used in [<sup>18</sup>F]FPS-ZM1 biodistribution and blood sampling

### Radiosynthesis

[^18^F]SDM-8 (0.37 GBq/ml) [25] (**SFig. 1**) and [^18^F]flumazenil (0.37 GBq/ml) [28] (**SFig. 2**) were synthesized as described earlier. [^18^F]PM-PBB3 (1.48 GBq/ml) was synthesized from an automatic synthesis module and kit provided by APRINDIA therapeutics (Suzhou, China) [36]. [^18^F]florbetapir (0.56 GBq/ml) was radiosynthesized from its precursor in a fully automated procedure suitable for routine clinical application [37, 38]. [^18^F]FPS-ZM1 (N-benzyl-4-chloro-N-cyclohexylbenzamide, 30 mCi/mL) was radiosynthesized from its precursor, as described in detail in the Supplementary file (**SFigs. 3**) [30]. The identities of the standard precursor [^18^F]FPS-ZM1 were confirmed by using mass spectrometry (MS), nuclear magnetic resonance (NMR) spectroscopy and high-performance liquid chromatography (HPLC, **SFigs. 4-7, STables 1, 2)**. The identities of the final products were confirmed by comparison with the HPLC retention time of the nonradioactive reference compound by coinjection using a Luna 5 μm C18(2) 100 Å (250 mm × 4.6 mm) column (Phenomenex) using acetonitrile and water (60:40) solvent with a 1.0 mL/min flow rate. Radiochemical purity > 95% was achieved for all aforementioned tracers.

### MicroPET

PET experiments using [^18^F]SDM-8, [^18^F]flumazenil, [^18^F]florbetapir, [^18^F]PM-PBB3 and [^18^F]FPS-ZM1 were performed using a Siemens Inveon PET/CT system (Siemens Medical Solutions, Knoxville, United States) [39]. Prior to the scans, rats were anaesthetized using isoflurane (1.5%) in medical oxygen (0.3-0.5 L/min) at room temperature with an isoflurane vaporizer (Molecular Imaging Products Company, USA). The rats were positioned in a spread-supine position on the heated imaging bed and subjected to inhalation of the anaesthetic during the PET/computed tomography (CT) procedure. The temperature of the rats was monitored. A single dose of tracers (∼0.37 MBq/g body weight, 0.2–0.5 mL) was injected into the animals through the tail vein under isoflurane anaesthesia. Static PET/CT imaging was obtained for 10 min post intravenous administration, depending on the tracer [^18^F]SDM-8 at 60 min [25], [^18^F]flumazenil at 20 min, [^18^F]FPS-ZM1 at 90 min, [^18^F]florbetapir at 50 min and [^18^F]PM-PBB3 at 90 min [38]. PET/CT images were reconstructed using the ordered subsets expectation maximization 3D algorithm (OSEM3D), with 2 iterations, image zoom 1, matrix size of 128 × 128 × 159 and a voxel size of 0.815 mm × 0.815 mm × 0.796 mm. Data were reviewed using Inveon Research Workplace (IRW) software (Siemens). Attenuation corrections derived from hybrid CT data were applied.

### Biodistribution study

For evaluation of the biodistribution of [^18^F]FPS-ZM1, 54 mice (aged 1.5 months, 27 M, 27 F) were intravenously injected with [^18^F]FPS-ZM1 (140 μCi) via the tail vein. Six animals (3 M, 3 F) were euthanized at each time point: 2 min, 5 min, 10 min, 15 min, 30 min, 45 min, 1 h, 3 h, and 4 h post injection. The organs and brain regions of the mice were dissected, weighed and analysed for radioactivity using a gamma counter (**STable. 3**). Blood sampling was performed in six mice (3 M, 3F) at 2 min, 5 min, 10 min, 15 min, 30 min, 45 min, 1 h, 2 h, 3 h, and 4 h post injection.

### Imaging Data Analysis

Images were processed and analysed using PMOD 4.4 software (PMOD Technologies Ltd., Zurich, Switzerland). Radioactivity is presented as the standardized uptake value (SUV) (decay-corrected radioactivity per cm^3^ divided by the injected dose per gram body weight) [40]. The time—activity curves were deduced from specific volumes of interest that were defined based on a rat MRI T_2_-weighted image template (W. Schiffer). Brain regional SUVR was calculated using the cerebellum (CB) as the reference region. The mask was applied for signals outside the brain volumes of interest for illustration (**SFig. 8**).

### Proteomics profiling and pathway analysis

After in vivo imaging, high-throughput quantitative proteomics analysis was performed on the hippocampal samples from two 12-month-old ZDF and two age-matched SD rats using tandem mass tag (TMT) labelling (**STable 2, 3**). Hippocampal rat brain tissues were collected and homogenized on ice for 10 min and lysis buffer containing 4% SDS, 7 mM urea, 30 mM N-2-hydroxyethylpiperazine-N’-2-ethanesulfonic acid HEPES, 1 mM phenylmethylsulfonyl fluoride, 2 mM Ethylenediaminetetraacetic acid, 2 mM thiourea, 10 mM DL-dithiothreitol (DTT), and 1× protease inhibitor. The supernatant was centrifuged at 10000 × g for 30 min at 4°C. The supernatant was then incubated with 100 mM tetraethylammonium bromide (TEAB) and 10 mM DTT for 60 min at 55°C, and then 35 mM iodoacetamide was added for 60 min in the dark and 5 times acetonitrile (v/v) at -20°C for 3 h. The sample was centrifuged at 20000 × g for 30 min at 4°C. The remaining sample was then incubated twice with 1 mL of 50% acetonitrile at -20°C for 3 h and then centrifuged at 20000 × g for 30 min at 4°C. Then, 0.1 mL of 100 mM TEAB was added to the protein precipitate and mixed. The peptide/protein concentration was determined using the BCA protein assay. The protein precipitate was added to 100 μL of 100 mM TEAB, 1 mg/ml trypsin, and 1.0 mg enzyme/100 mg protein and incubated at 37°C for 4 h. Then, trypsin 1.0 mg was added, followed by incubation at 37°C for 12 h. The peptide solution was centrifuged at 5,000 g and dried into powder using a freeze dryer. Detection of enzymatic hydrolysis efficiency was performed using liquid chromatographyDmass spectrometry. The peptide/protein concentration in the supernatant was determined by BCA protein assay.

TMT labelling of peptides was performed according to the instructions of the TMT 6-plex kit. Peptides were analysed using a liquid chromatography system. Extracted spectra from Proteome Discoverer (2.4.0.305) were searched with Sequest HT. Quantitative analysis of the spectra resulting from the first step was performed using Proteome Discoverer. The false positive rate (FDR) was set at ≤ 1% for both protein and peptide levels (P value <0.05; fold change >1.2).

In addition, a comprehensive bioinformatics analysis was used to investigate the differentially expressed proteins and enriched signaling pathways as described earlier [41-43]. We analysed the differentially expressed genes (DEPs) by using principal component analysis (PCA), volcano plot analysis, subcellular localization analysis, Cluster of Orthologous Groups of proteins (COG) analysis, kinase analysis, proteomic and peptide assays, (Kyoto Encyclopedia of Genes and Genomes, KEGG) signaling pathways, modification site assay, differential gene expression analysis, and gene ontology (GO) analysis. KEGG signaling pathway analysis was performed using the database for *Rattus norvegicus* (rat) (www.kegg.jp/kegg/pathway.html). GO analysis was performed using the GO database (http://www.geneontology.org/) with annotations based on the UniProt database (https://www.uniprot.org/taxonomy/10116). ProteinDprotein interaction (PPI) network analysis was performed for the DEPs using the STRING database for *Rattus norvegicus* (rat) (v11, stringdb.org).

### Statistics

Two-way ANOVA with Sidak post hoc analysis was used for comparisons between groups (GraphPad Prism 9.0, CA, USA). Pearson correlation analysis was used for the analysis of the association between the regional SUVR of different tracers. Significance was defined as p < 0.05. Data are shown as the mean ± standard deviation.

## Results

### No detectable PET changes in amyloid-beta or tau levels in ZDF rat brains

We observed comparable levels of [^18^F]florbetapir (ZDF rat, n=15; SD rat, n=8, **Figs. 1a-c**) and [^18^F]PM-PBB3 (ZDF rat, n=7; SD rat, n=4, **Figs. 1d-f**) in the brains of 12-month-old ZDF vs SD rats. This suggested that there was not sufficient accumulation of amyloid-beta and tau aggregates in the brains of ZDF rats at 12 months of age.

**Fig. 1.**
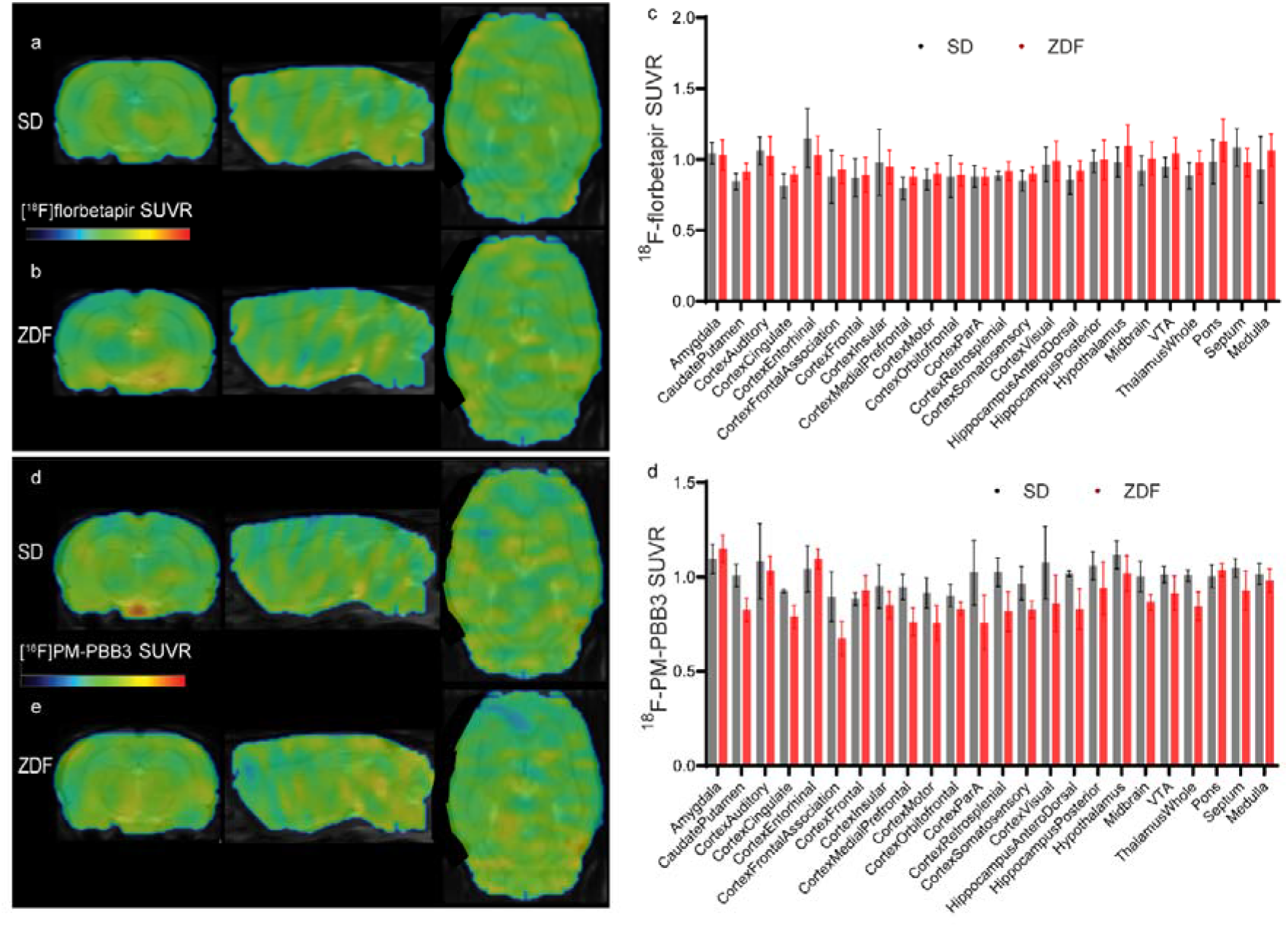
No detectable amyloid-beta and tau changes by [^18^F]florbetapir and [^18^F]PM-PBB3 brain uptake in 12-month-old ZDF compared to age-matched SD rats. (a-b) [^18^F]florbetapir and (d-e) [^18^F]PM-PBB3 SUVR images (scale 0-2.2). (c, f) Quantification of [^18^F]florbetapir SUVR in the brain (ZDF rat, n=15; SD rat, n=8) and [^18^F]PM-PBB3 (ZDF rat, n=7; SD rat, n=4).

### Reduced regional levels of SV2A and GABA_A_ receptors in ZDF rat brains

The level of SV2A revealed by [^18^F]SDM-8 was reduced in the parietal cortex in 12-month-old ZDF rats compared with age-matched SD rats (p = 0.0087, ZDF rat, n=3; SD rat, n=3, **Figs. 2a-c**). [^18^F]flumazenil SUVR, indicative of GABA_A_ receptor levels, was reduced by approximately 30% in the frontal cortex (p = 0.0015), approximately 25% in the somatosensory cortex (p = 0.0244), and approximately 30% in the hippocampus (anterior dorsal p = 0.0092, posterior p = 0.0417) of 12-month-old ZDF rats compared to age-matched SD rats (ZDF rat, n=3; SD rat, n=3, **Figs. 2d-f**).

**Fig. 2.**
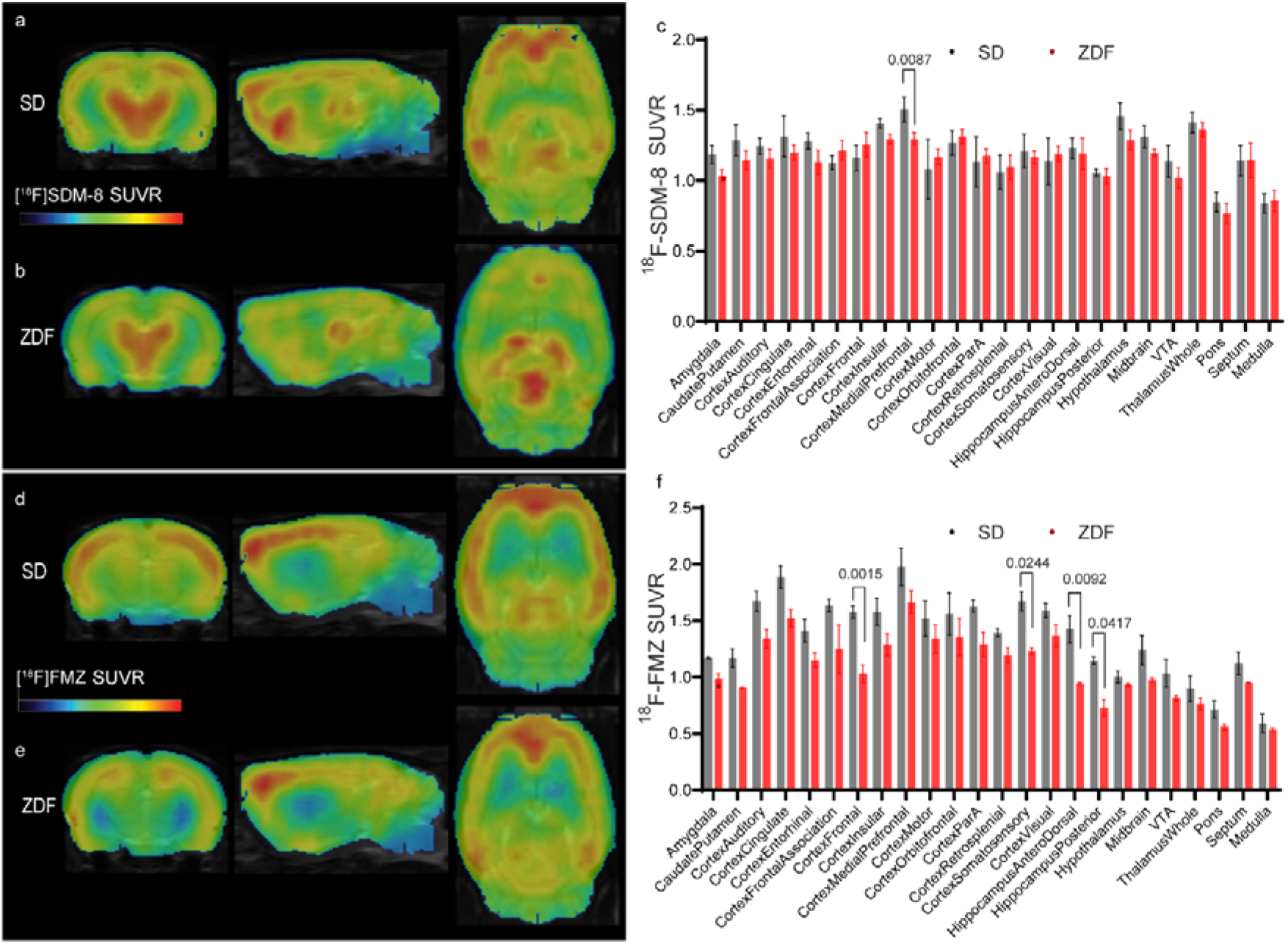
Reduced regional levels of SV2A and GABA_A_ receptors in the brains of 12-month-old ZDF rats compared to those of age-matched SD rats. (a-b) [^18^F]SDM-8 and (d-e) [^18^F]flumazenil (FMZ) SUVR images (scale 0-2.2). (c, f) Quantification of [^18^F]SDM-8 SUVR in the brain (ZDF rat, n=5; SD rat, n=4) and [^18^F]FMZ (ZDF rat, n=3; SD rat, n=3).

### Proteomics profiling and pathway analysis

A total of 309 DEPs were detected in the hippocampus of 12-month-old ZDF rats compared to SD rats, mainly annotated as cytoplasmic (38.57%) and periplasmic (34.08%) (**Fig. 3b**). Downregulated proteins, including the synaptic-related proteins Dbnl, Syn1, Syn2, and Map2, were detected (**Fig. 3a**). The DEPs were mostly associated with signal transduction mechanisms and enriched in synapse and neuron signaling (**Figs. 3c, d**). Enrichment analysis of the DEPs indicated enrichment in the MAPK and cAMP insulin secretion-related pathways, endocrine and other factor-regulated calcium reabsorption, etc. (**Fig. 4a**), and the proteinDprotein interaction network with the proteins shown as nodes (**Figs. 4b, c**).

**Fig. 3.**
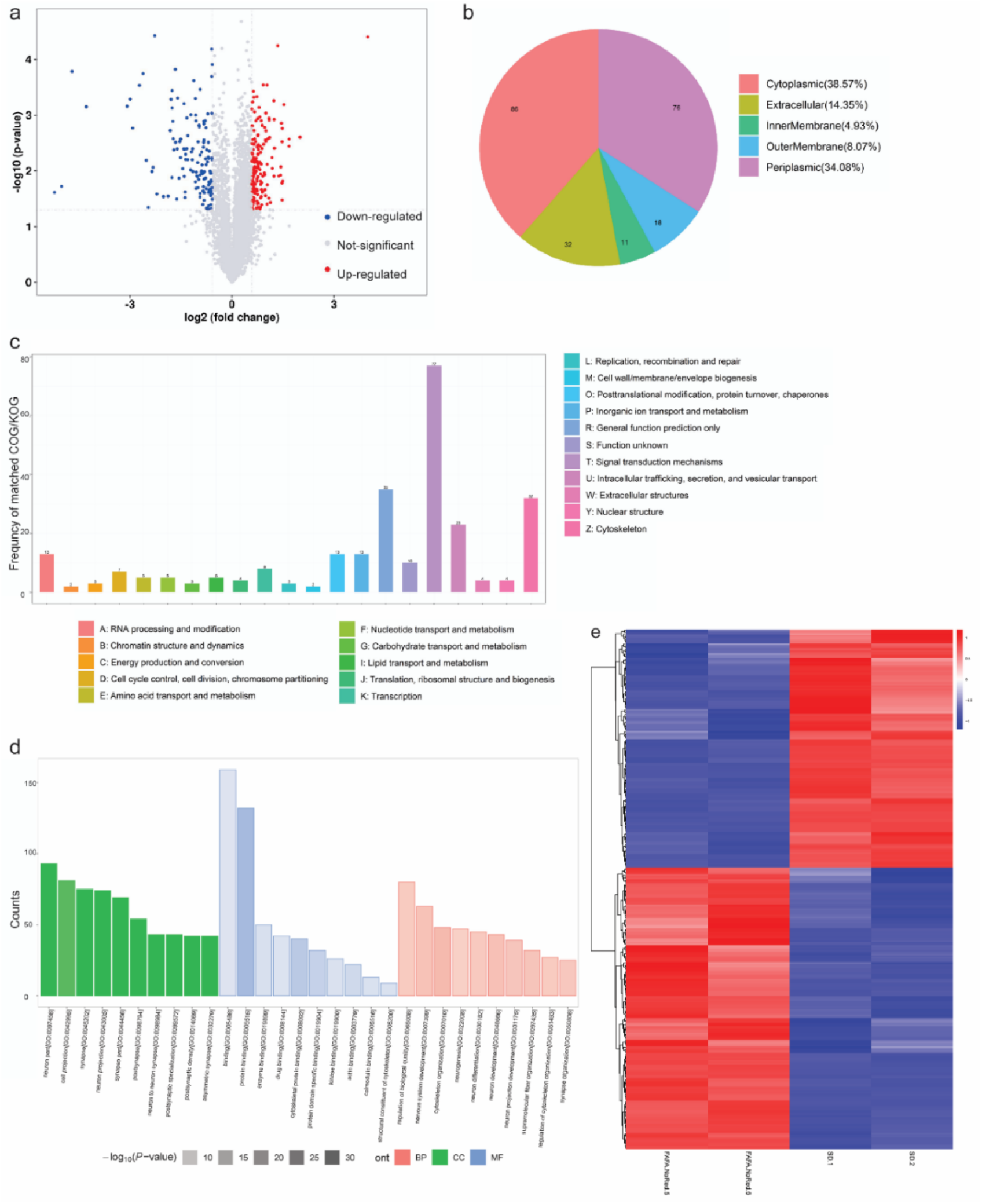
Profiling of differentially expressed proteins (DEPs) in the hippocampus of 12-month-old ZDF and SD rats. (a) Volcano plots showing log2-fold-change (x-axis) and -log10 p value (y-axis) for all quantified proteins. (b) Location of DEPs. (c) The number of matched genes assigned in clusters of orthologous groups (COGs) and eukaryotic orthologous groups (KOGs). (d) Gene Ontology (GO) enrichment analysis of biological process (BP), cellular component (CC), and molecular function (MF) terms. The number on the x-axis indicates the enriched count of DEPs. (e) Heatmap for DEPs.

**Fig. 4.**
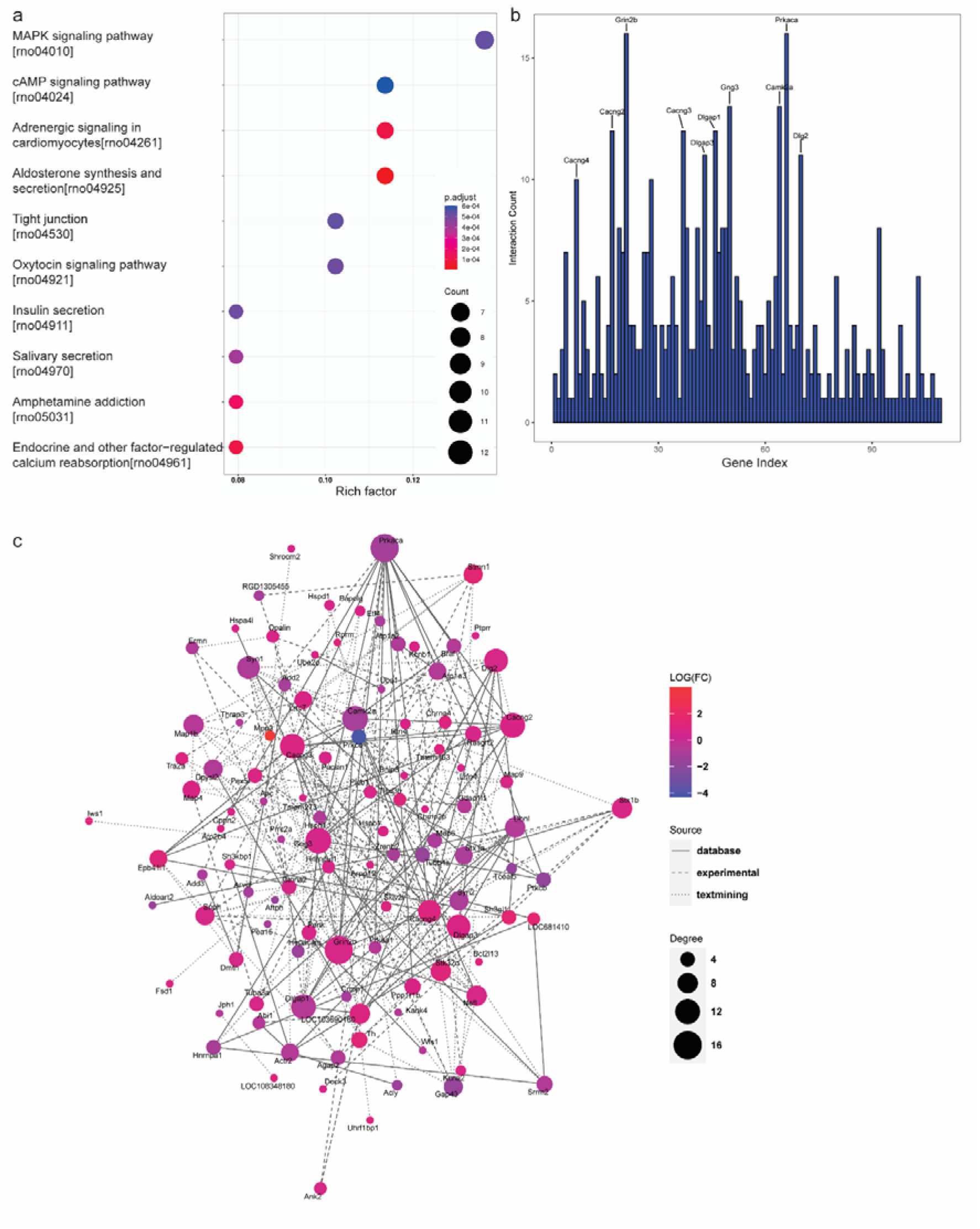
Kyoto Encyclopedia of Genes and Genomes (KEGG) pathway analysis of differentially expressed proteins (DEPs) enriched in pathways. (a) Enrichment analysis showing the top 10 enriched pathway terms. (b, c) ProteinDprotein interaction network analysis. Histogram of interaction (b). The proteins present in the networks are shown as nodes (c).

### Large variation in [^18^F]FPS-ZM1 uptake in the brains of ZDF rats

We first assessed the biodistribution of [^18^F]FPS-ZM1 in mice. High variation in [^18^F]FPS-ZM1 levels in the various brain regions and high liver uptake were observed by ex vivo biodistribution in C57BL/6J mice (**SFig. 9a-c**). The brain uptake of [^18^F]FPS-ZM1 was lower in the brain than in the spinal cord and in the upper trunk in 12-month-old ZDF rats (**SFigs. 9d-g**). Large variation in the regional distribution of RAGE in the brains of 12-month-old ZDF rats by PET using [^18^F]FPS-ZM1 was also observed.

## Discussion

Thus far, few imaging studies have been reported on alterations in the central nervous system of T2DM models, including [^18^F]FP-CMT for dopamine innervation[44] and [^18^F]FDG for metabolism. Cognitive impairment, cortical thinning, and hippocampal synaptic reorganization have been observed in T2DM rats [6]. Here, we demonstrated synaptic and GABAergic impairments without detectable changes in the levels of Aβ and tau by PET in the brains of 12-month-old ZDF rats compared to SD rats. To our knowledge, this is the first study that visualized these targets noninvasively in the brains of T2DM animal models.

One possibility for the lack of differences in the levels of [^18^F]florbetapir and [^18^F]PM-PBB3 uptake could be that the accumulation levels of Aβ and tau deposits have not reached the threshold detectable by using PET or that the aggregates are not mature enough to harbor sufficient fibrillar structures that the β-sheet binding imaging tracers recognize [45, 46]. This is reflected partly in previous studies using immunohistochemical staining showing relatively low insoluble Aβ, Aβ plaques, and tau deposits in the brains of T2DM rats [9-11] compared to those in animal models of amyloidosis [47, 48] and tauopathy [49, 50].

Our finding of reduced cortical levels of SV2A measured by [^18^F]SDM-8 in the brains of aged ZDF rats compared to SD rats corroborated our findings of downregulation of the synaptic-related proteins Dbnl, Syn1, Syn2, and Map2 and synaptic-related pathways in hippocampal tissues from 12-month-old ZDF rats compared to SD rats. This is in line with earlier biochemical studies [5-7], and proteomic studies in the hippocampus of ZDF rats identified reductions in the levels of the metalloproteins apolipoprotein C, myelin binding protein and apolipoprotein A-I as well as the synaptic proteins Mapk1, OMG, Gng12, Dbnl, sv2a, and Stx12 [41-43].

Our observation of the reduced [^18^F]flumazenil measure of GABA_A_ receptors in the cortex and hippocampus of aged ZDF rats compared to SD rats is in line with the known alterations of GABA_A_ receptors in the brain of ZDF rat models [4]. Given the important role of the GABAergic system in diabetic cognitive dysfunction, blood sugar control and energy homeostasis [51], the GABAergic system has been an important drug target for T2DM. Therefore, [^18^F]flumazenil might be useful in further investigation and monitoring of therapeutics targeting GABAergic impairment in T2DM.

Autoradiography using [^11^C]FPS-ZM1 in aged Tg2576 amyloidosis mice [31], as well as [^18^F]InRAGER (intracellular domain of RAGE) in a lipopolysaccharide model, has been reported [52]. However, the relatively suboptimal affinity (15 nM), high lipophilicity (CLogP: 5.25), and rapid metabolism of [^18^F]FPS-ZM1 limited its in vivo performance [30]. Our results of the biodistribution and pharmacokinetics and in vivo [^18^F]FPS-ZM1 in ZDF rats are in line with earlier observations [52]. Further optimization of tracers targeting RAGE is needed for in vivo application.

There are several limitations of this study. First, only male ZDF and SD rats at 12-months old were investigated. Further imaging studies including both male and female rats at different ages and stages of disease are needed to assess the. In addition, we did not include immunohistochemical staining to validate the in vivo imaging results at postmortem.

In summary, we demonstrated in vivo regional reductions in SV2A and GABA_A_ receptors in the brains of aged ZDF rats without detectable Aβ and tau accumulation. Further study using a balanced and longitudinal design will provide further insights into the dynamics of SV2A and GABA_A_ receptor alterations in the brains of T2DM rats, which will be useful in understanding cognitive impairment in T2DM and monitoring treatment.

## Authors’ contributions

The study was designed by KY and RN. KY and QH performed radiolabelling, and KY performed HPLC and microPET. RN performed the microPET analysis. CZ, JD, and JX prepared tracers. KY and RN wrote the first draft. All authors contributed to the revision of the manuscript and approved the final manuscript.

## Data availability

The datasets generated and/or analysed during the current study are available from the corresponding author upon reasonable request.

## Ethics approval

The PET imaging and experimental protocol was approved by the Institutional Animal Care and Ethics Committee of Fudan University and performed in accordance with the National Research Council’s Guide for the Care and Use of Laboratory Animals.

