## Supplementary files for "Reduced SV2A and GABA_A_ receptor levels in the brains of type 2 diabetic rats revealed by [^18^F]SDM-8 and [^18^F]flumazenil PET"

**STable 1 Instrument used in the synthesis and identification of chemicals and proteomics analysis;**

| **Instrument used in radiosynthesis** | **No** | **Company** |
| --- | --- | --- |
| Dose Calibrator  High-performance liquid Chromatography (HPLC) | ATOMLAB^TM^ 500  Prep2100, 1500 | BIODEX  Alltech |
| Online radiation detector | FC1000 | Bioscan |
| Nuclear magnetic resonance | AVANCE III 400 | Bruker BioSpin AG |
| Mass spectrometry | Q Exactive Plus spectrometer | Thermo Scientific |
| Thin-layer chromatography | MiniScan | Bioscan |
| Multi Purpose Synthesizer | CFN-MPS20 | Sumitomo Heavy Industries |
| Radio-immunity gamma-counter | GC-1200 | KeDa Innovation Co., Ltd. |
| **Instruments used in proteomics** | **No** | **Company** |
| H-Class ultrahigh pressure liquid chromatography |  | Waters |
| Ultrahigh-pressure nanoscale liquid chromatography | EASY-nLC 1000 | Thermo Scientific |
| Orbitrap Fusion mass spectrometer |  | Thermo Scientific |
| Refrigerated tabletop centrifuge | 5417R | Eppendorf |
| Palm centrifuge | D1008 | Sero Czech SCILOGEX |
| Ultrasonic Crusher | Q125 | Qsonica |
| Freeze dryer | RVC 2-25 CDplus, CT 02-50 cold trap | Christ, Germany |
| Enzyme analyser | MB-102 | Hangzhou BORI Technology Co., Ltd. |

**STable 2 Materials used in the proteomic analysis**

| **Item** | **Comapny** |
| --- | --- |
| Urea, analytically pure | GibcoBRL |
| Sequencing grade Trypsin | Promega |
| BCA protein quantification kit | Fisher Scientific |
| Trifluoroacetic acid (TFA) | Sigma |
| Ammonium formate | Sigma |
| TMT6 labelling kit | Fisher Scientific |
| PMSF, ultra pure grade | Amesco |
| Ethylenediaminetetraacetic acid (EDTA), ultra pure grade | Amesco |
| 4-hydroxyethylpiperazineethanesulfonic acid (HEPES) | Sigma |
| Protease inhibitors | Roche, Switzerland |
| Dithiothreitol (DTT), chemically pure | Promega |
| Iodoacetamide (IAA), chemically pure | Promega |
| Sodium dodecyl sulfate, chemically pure | Sigma |
| Ethanol, mass spectrometry pure | Fisher Scientific |
| Formic acid, mass spectrometry pure | Fisher Scientific |
| Acetonitrile, mass spectrometry pure | Fisher Scientific |
| Acetone, mass spectrometry pure | Fisher Scientific |
| Triethylammonium bicarbonate (TEAB) | Santa Cruz |
| 25% Ammonia | Santa Cruz |
| sep-Pak C18 desalting column, 1 cc (100 mg) | Waters Corporation |
| High-Select™ Fe-NTA Phosphopeptide Enrichment Kit, Catalog Number A32992 | Fisher Scientific |
| High pH reversed-phase column, Acquity UPLC®BEH C18 1.7 µm, 2.1×50 mm | Waters Corp. |
| Nanoscale peptide analysis column, Acclaim PepMap C18, 75 µm×250 mm | Thermo Scientific |

**STable 3 Stability of [^18^F]FPS-ZM1 in buffer and in plasma**

| **Time (h)** | **Purity in buffer %** | **Purity in plasma%** |
| --- | --- | --- |
| 0 | 100 | 100 |
| 0.5 | 100 | 100 |
| 1 | 100 | 100 |
| 1.5 | 100 | 100 |
| 2 | 100 | 100 |
| 3 | 100 | 100 |
| 4 | 100 | 100 |
| 5 | 100 | 100 |
| 6 | 93 | 55 |

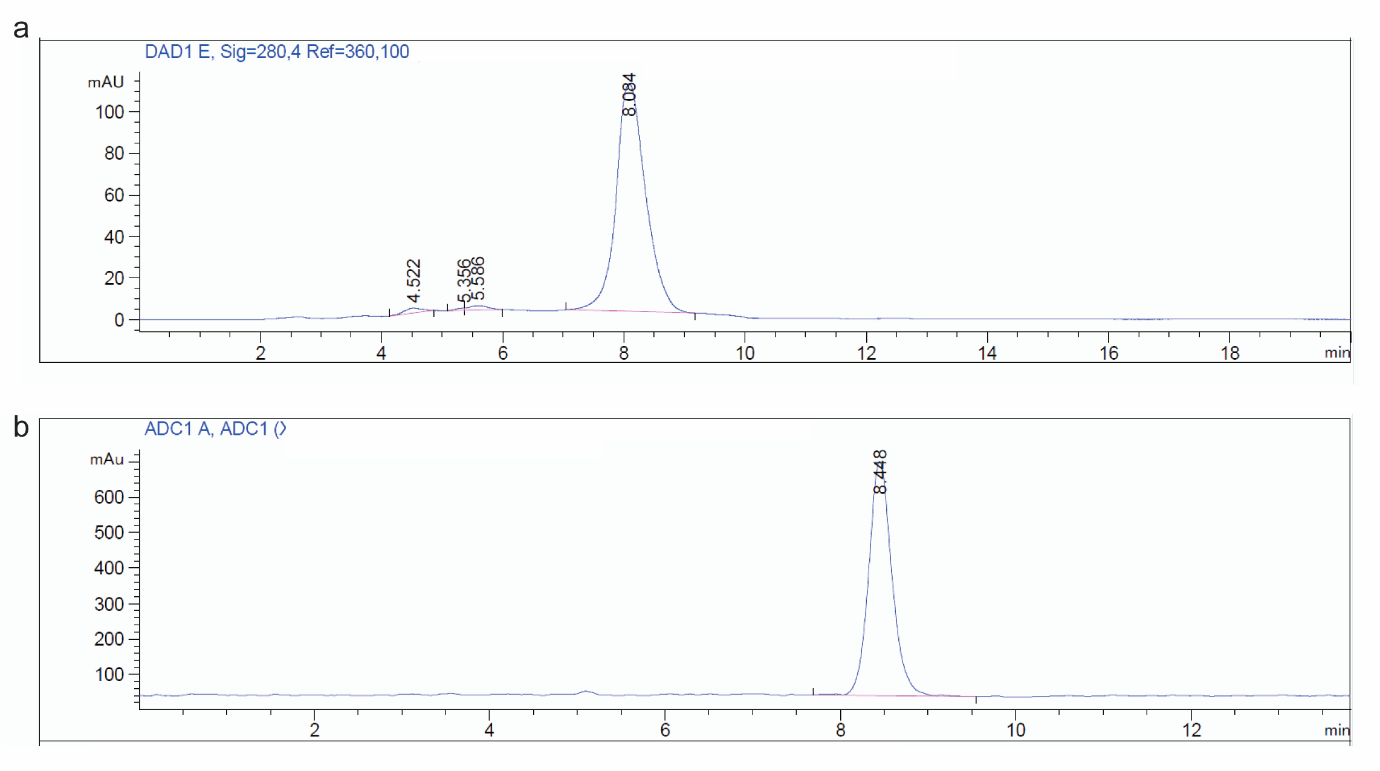

**Supplementary Fig 1. HPLC analysis of synthesized [^18^F]SDM-8**. (a) HPLC chromatogram for standard, (b) HPLC chromatogram for synthesized [^18^F]SDM-8.

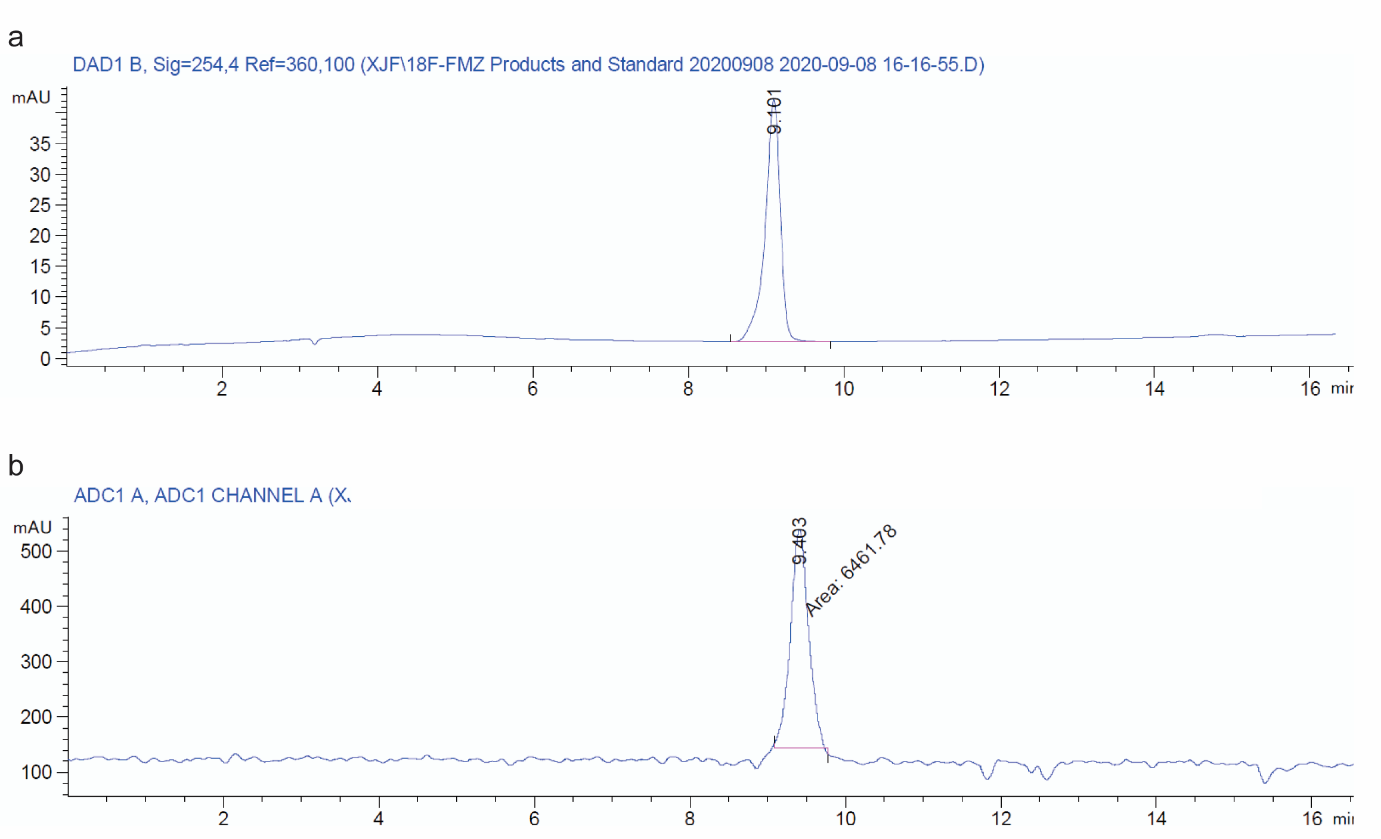

**Supplementary Fig 2. HPLC analysis of synthesized [^18^F]FMZ**. (a) HPLC chromatogram for standard, (b) HPLC chromatogram for synthesized [^18^F]FMZ.

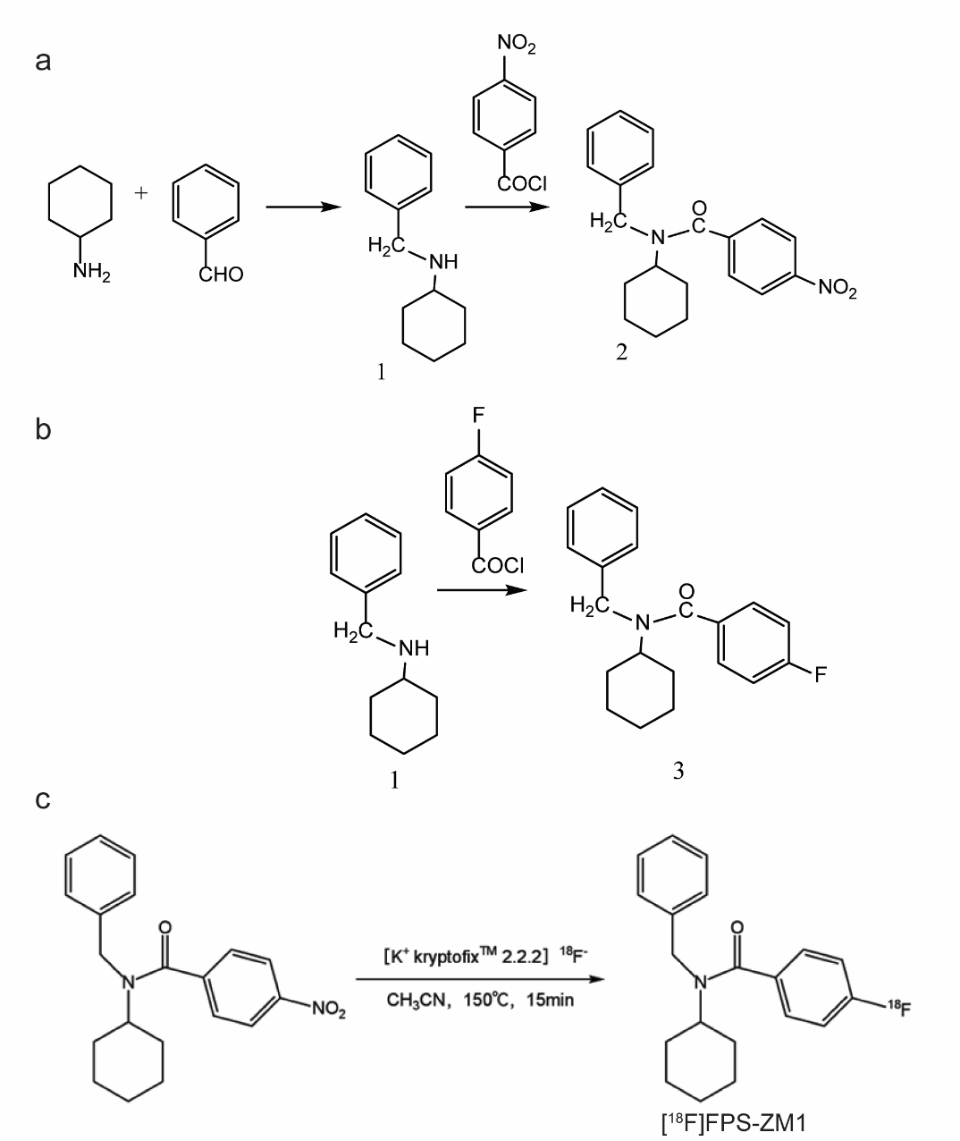

**Supplementary Fig 3. Synthesis of precursor, standard and radiolabelling for [^18^F]FPS-ZM1.** Synthesis of (a) precursor NO2-FPS-ZM1, (b) standard F-FPS-ZM1, and (c) radiolabelling of [^18^F]FPS-ZM1

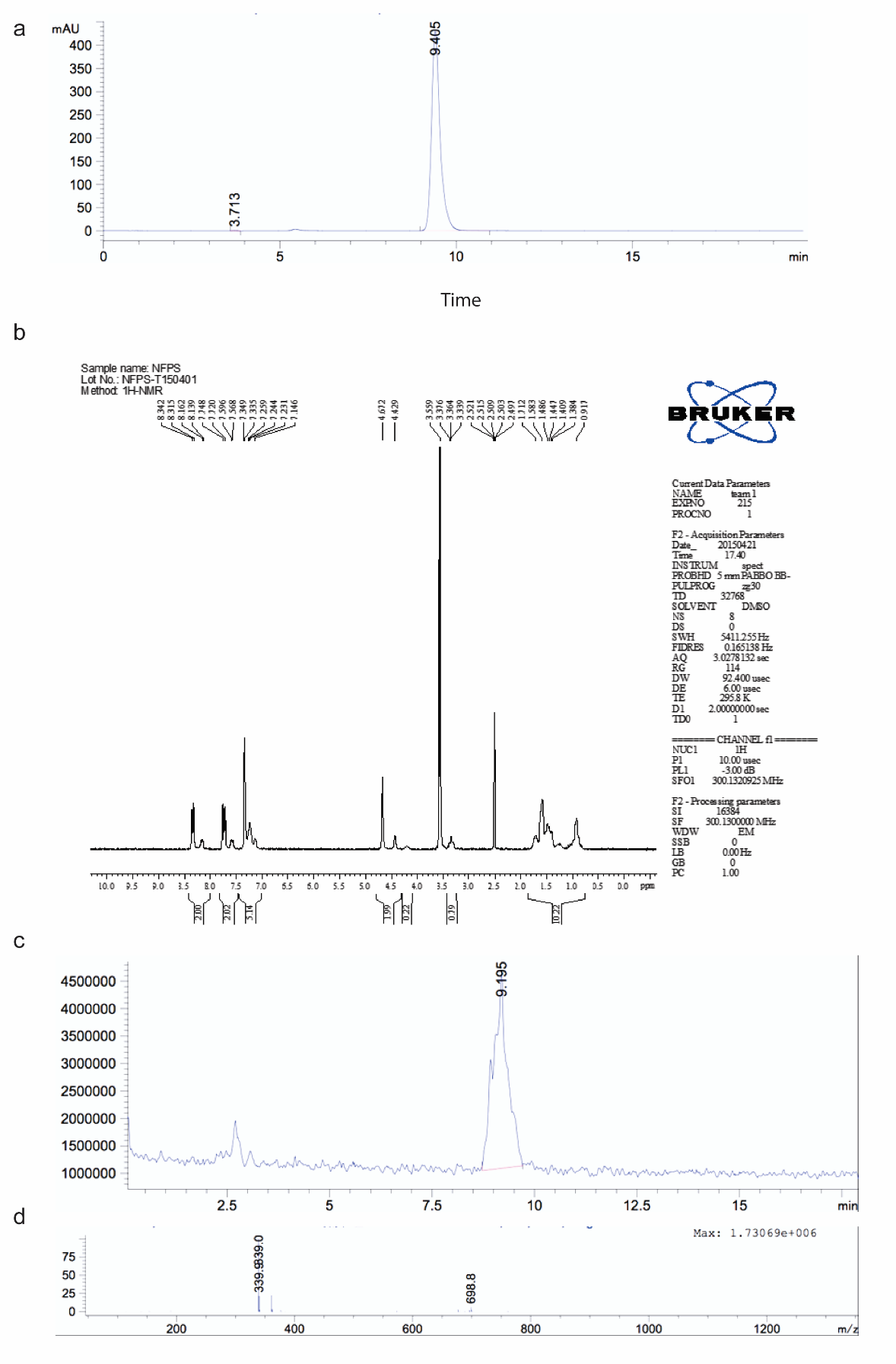

**Supplementary Fig 4. Identification of synthesized precursor NO2-FPS-ZM1.** (a) HPLC (b) ^1^H nuclear magnetic resonance (300 MHz, d-DMSO) δ 8.13-8.34(m,2H), 7.56-7.75(m, 2H), 7.14-7.35(m, 5H), 4.43-4.67(m, 2H), 3.33-3.38(m, 1H), 0.91-1.71(m, 10H); LC‒MS: calculated for C20H22N2O3，338.16；found [M+H] 339.0. (c, d) Mass spectrometry

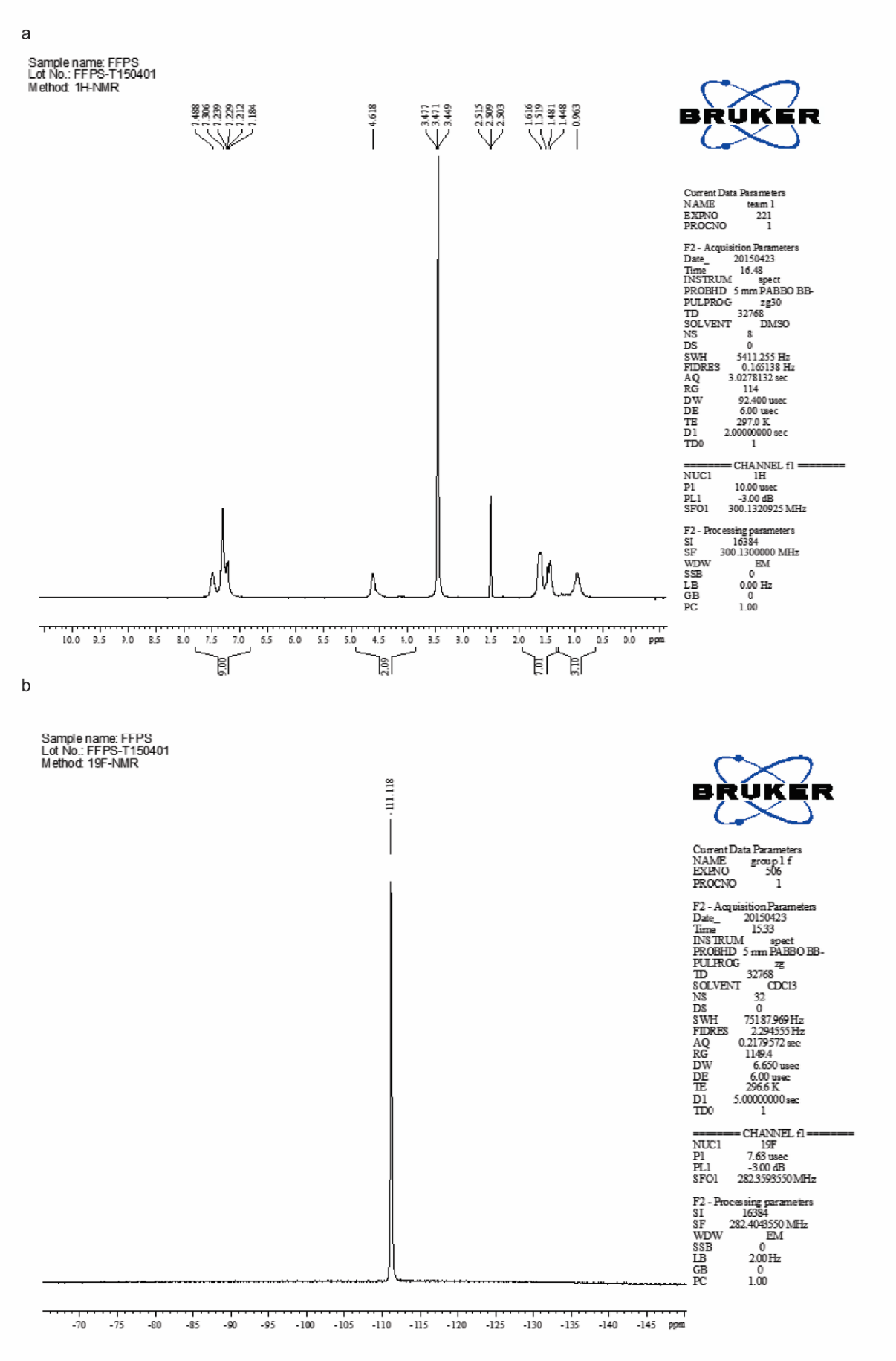

**Supplementary Fig 5. F-FPS-ZM1 standard identification. (a)** 1H nuclear magnetic resonance (300 MHz, d-DMSO) δ 7.13-7.49 (m, 9H), 4.48-4.62 (m, 2H), 3.28-3.45 (m, 1H), 0.96-1.61 (m, 10H); LC‒MS: calculated for C20H22FNO, 311.17; found [M+H] 312.0. (b) 19F NMR (300 MHz, CDCl3) δ -111.118.

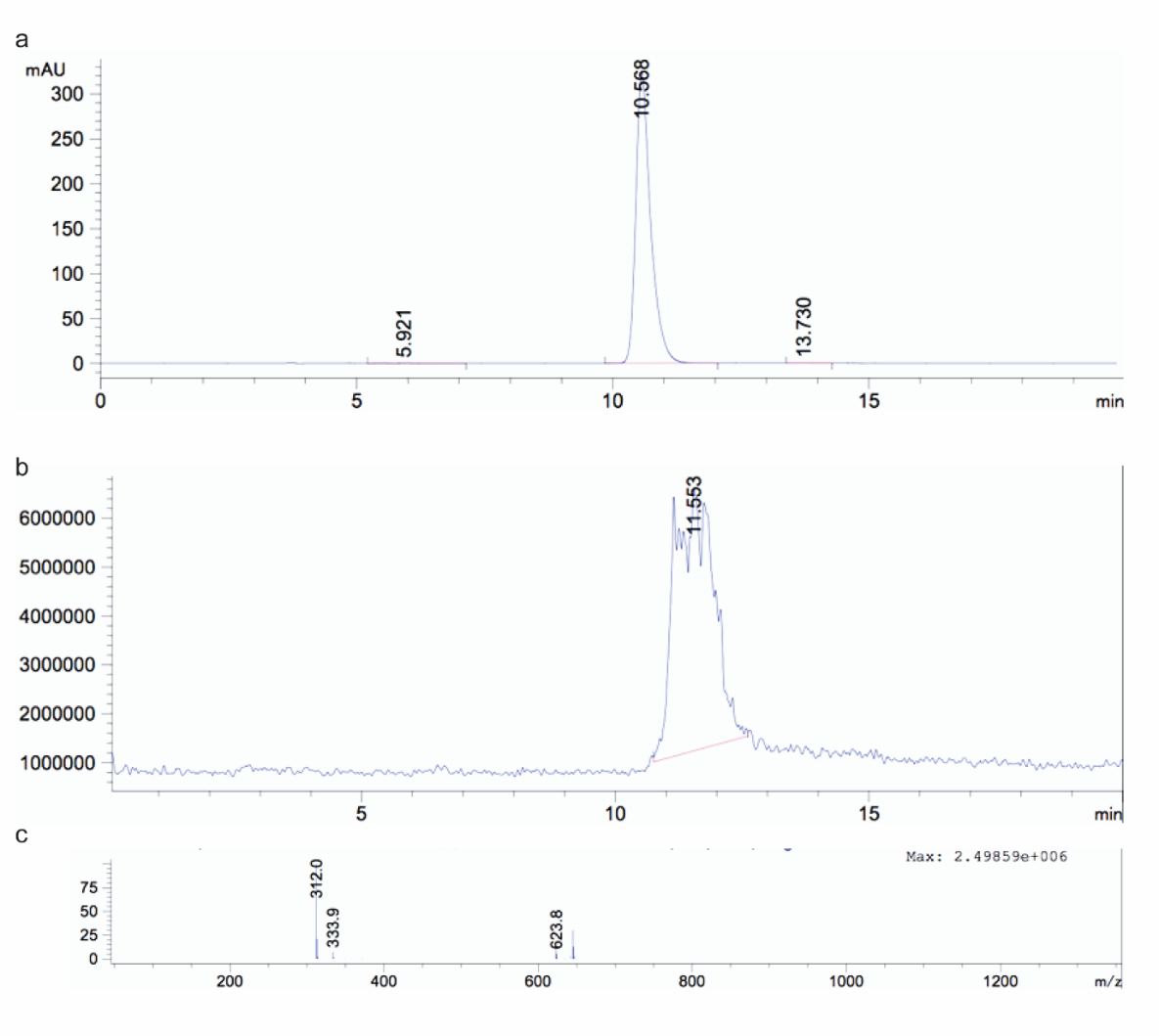

**Supplementary Fig 6. [^18^F]FPS-ZM1 standard identification. F-FPS-ZM1** (a) HPLC (b, c) Mass spectrometry

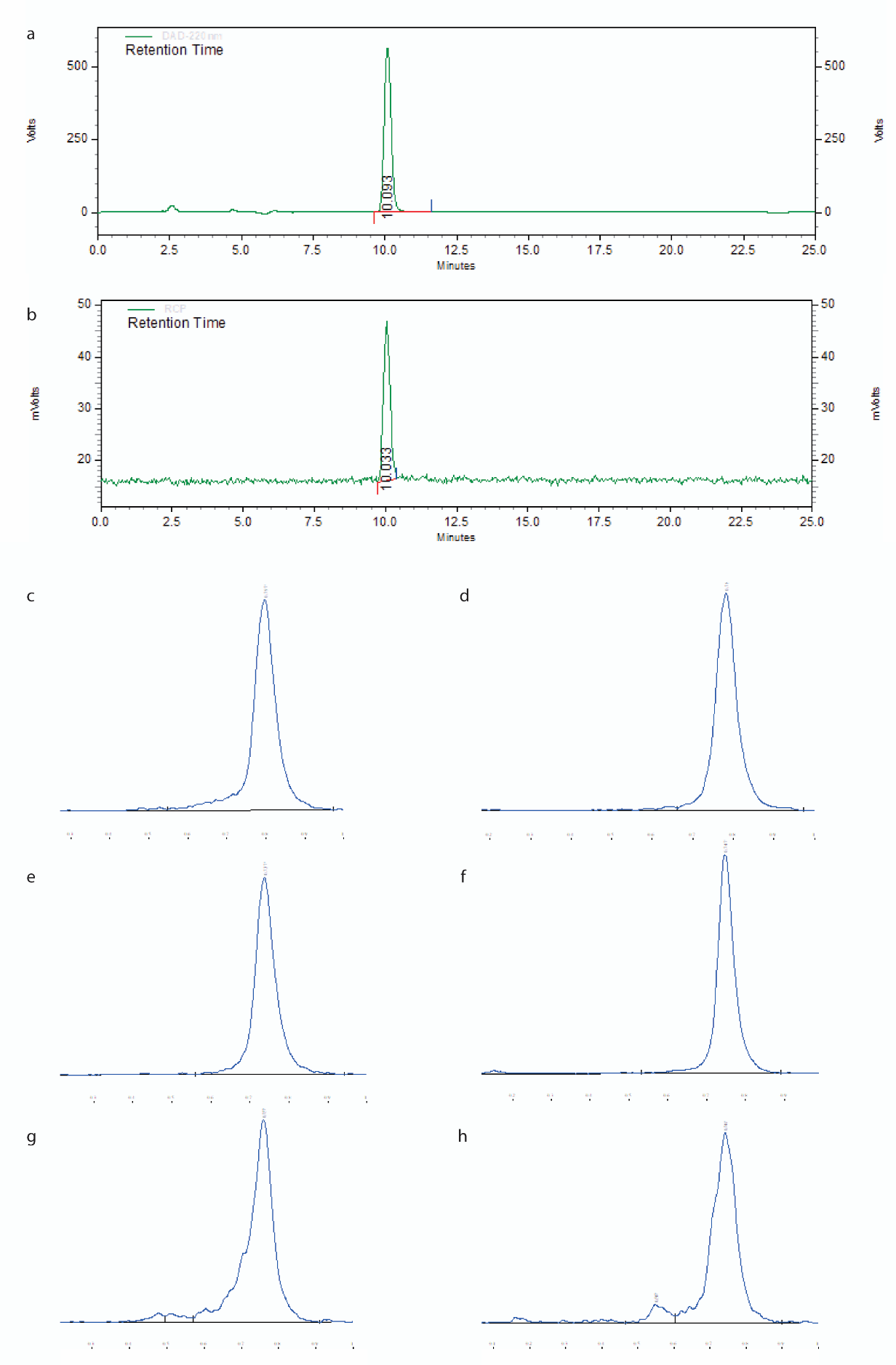

**Supplementary Fig 7. HPLC and thin-layer chromatography analysis of synthesized [^18^F]FPS-ZM1**. (a) HPLC chromatogram for standard, (b) HPLC chromatogram for synthesized [^18^F]FPS-ZM1, and (c-h) thin-layer chromatography of synthesized [^18^F]FPS-ZM1 at 0, 1, 2, 3, 4, and 6 hours after radiolabelling.

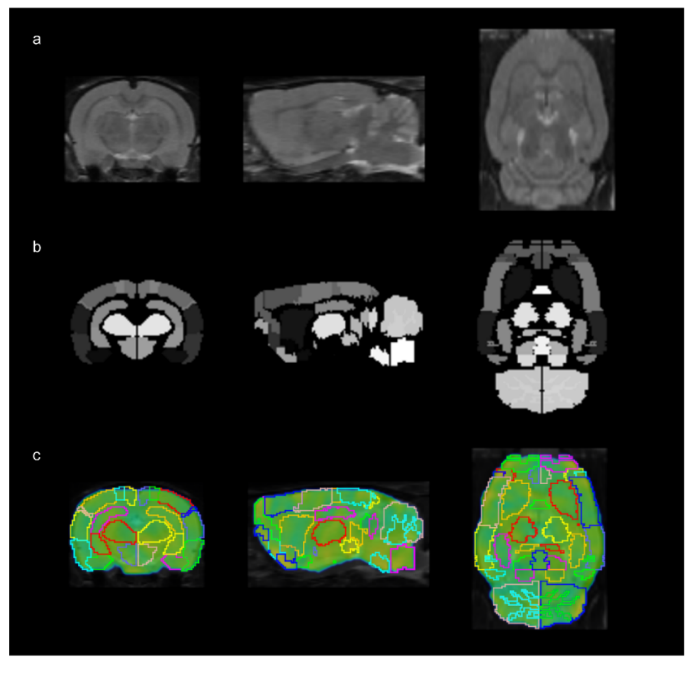

**SFig. 8 Volume-of-interest analysis of PET data in the rat brain.** (a) T_2_ MRI from the W. Schiffer rat brain template. (b) Color-coded brain region. (c) Overlay of VOI outline on PET images.

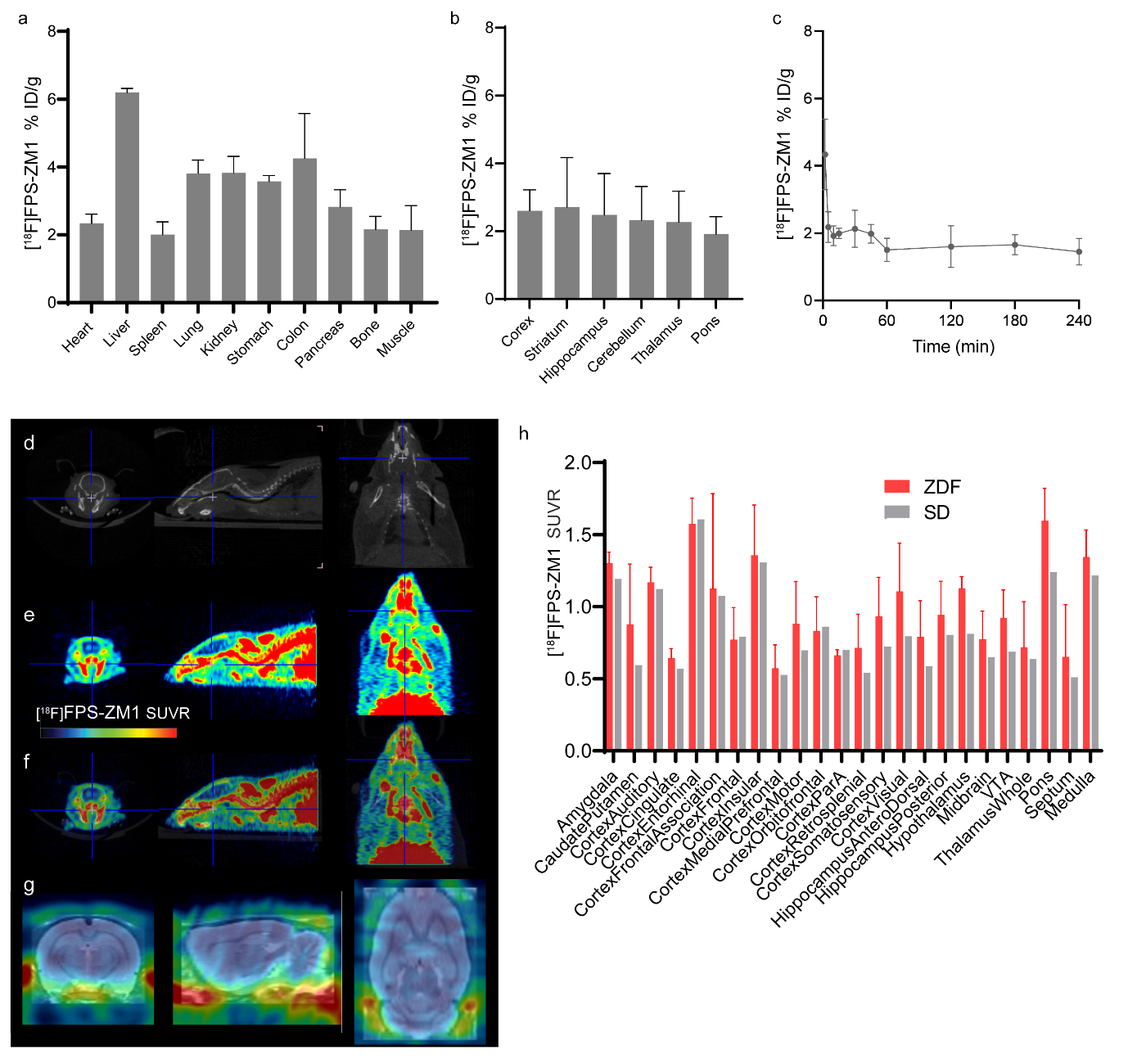
**SFig. 9** **[^18^F]FPS-ZM1 brain uptake in 12-month-old ZDF rats**. (a, b)Biodistribution in organs and brain regions (n=6/time point). (c)Time activity curve in the blood sampled post injection(n=6/time point). (d-g) Representative [^18^F]FPS-ZM1 PET in ZDF rat, CT(d), PE (e), PET/CT overlaid view(f), PET/MRI overlaid(g). SUVR scale 0-4.8. (i)Quantification of [^18^F]FPS-ZM1 SUVR in the brain(ZDF rat, n=5; SD rat, n=1).
